# The entopeduncular nucleus drives lateral habenula responses to negative but not positive or neutral affective stimuli

**DOI:** 10.1101/408963

**Authors:** Hao Li, Dominika Pullmann, Thomas C. Jhou

## Abstract

Lateral habenula (LHb) neurons are activated by negative motivational stimuli and play key roles in the pathophysiology of depression. Early reports indicated the possibility that rostral entopeduncular nucleus (rEPN) neurons drive these LHb responses, but this influence remains untested. We find that both rEPN and LHb neurons in rats exhibit similar activation/inhibition patterns after negative/positive motivational stimuli, but that the rEPN influence on LHb firing is surprisingly selective. Temporary rEPN inactivation decreases LHb basal and burst firing, and eliminates LHb responses to footshock-predictive cues occurring 40-100ms but not 10-30ms post-stimulus, nor on responses to positive/neutral motivational stimuli. Additionally, rEPN inactivation partially but not fully reduces LHb responses to signaled footshocks, while excitotoxic rEPN lesions only partially diminish footshock-induced cFos in the LHb and its rostromedial tegmental nucleus targets. Together, our findings indicate an important but selective role of the rEPN in driving LHb responses to motivational stimuli.

## Introduction

The lateral habenula (LHb) has been implicated in processing aversive events across many species (Jhou et al., 2013; Matsumoto and Hikosaka, 2008; Salas et al., 2010; Stopper and Floresco, 2014; Wang et al., 2017). Neurons in the LHb are activated by negative motivational stimuli, which in turn suppress dopamine (DA) firing via activation of GABAergic neurons in the rostromedial tegmental nucleus (RMTg) (Brown et al., 2017; Hong et al., 2011; Jhou et al., 2009a; Jhou et al., 2009b; Ji and Shepard, 2007). Dysregulation of this pathway can lead to cognitive disorders associated with abnormal processing of motivational stimuli including depression (Baker et al., 2016; Elmer et al., 2018; Lawson et al., 2017; Shumake and Gonzalez-Lima, 2003). Specifically, both rodent models and human patients with depression have shown that the pathophysiology of depression is highly associated with increased basal activities in LHb neurons and blunted LHb responses to negative events (Cui et al., 2018; Lawson et al., 2017; Li et al., 2013; Yang et al., 2018). Furthermore, synaptic potentiation onto LHb neurons in mice mimics core symptoms of depression, which can be alleviated with antidepressants (Li et al., 2011; Shabel et al., 2014), suggesting an important role of LHb afferents in mediating depressive symptoms. However, the identity of such afferents is unknown, as LHb neurons receive inputs from many brain regions (Yetnikoff et al., 2015), and the sources driving LHb basal activities and responses to negative stimuli remain incompletely characterized.

Studies in mice have indicated a close anatomical and functional relationship between LHb neurons and neurons in the rostral entopeduncular nucleus (rEPN), a region homologous to the internal globus pallidus (GPi) in primates, and sometimes referred to as the habenular-projecting cells of the globus pallidus (GPh). For example, the rEPN projections to the LHb can be excitatory and aversive in rodents, while primate GPi neurons respond to negative stimuli similarly to LHb neurons (Bromberg-Martin et al., 2010; Shabel et al., 2012; Stephenson-Jones et al., 2016). This evidence suggests that rEPN neurons would drive LHb activation to negative stimuli, but such an influence has not been directly tested. In fact, LHb neurons also receive glutamatergic inputs from several other regions which are also involved in motivational processing (Lecca et al., 2017; Root et al., 2014; Tooley et al., 2018), raising further questions about the relative role of the rEPN in negative encoding versus other LHb afferents.

In the present study, we compared neural activities induced by repeated footshocks in several LHb-projecting regions using cFos expression and retrograde tracing from the LHb. We further characterized response patterns of rEPN and LHb neurons to both reward and footshocks, as well as expectation of these stimuli using *in vivo* electrophysiology. Finally, we examined the effects of rEPN inactivation on LHb basal activities and responses to these stimuli, and tested effects of rEPN lesions on footshock-induced cFos in the LHb and RMTg. Taken together, our findings suggest a surprising complexity of both LHb firing patterns and the selectivity of rEPN influences on them.

## Results

### LHb-projecting neurons in the rEPN and the LH are significantly activated by repeated footshocks

In order to identify LHb afferents that respond to aversive stimuli, we used cFos as a proxy of neural activity and labeled LHb-projecting neurons by injecting the retrograde tracer cholera toxin subunit B (CTb) into the LHb (**Fig. 1A**). On the test day, animals were either given a series of footshocks (0.7mA, 30 times), or remained undisturbed in their home cage, before being sacrificed one hour later (n=7 per group). We calculated the proportion of CTb positive neurons expressing cFos in several LHb afferents including the ventral palladium (VP), lateral preoptic area (LPO), rEPN, lateral hypothalamus (LH), and ventral tegmental area (VTA). Compared with the control group, we found that footshock presentation excited over 80% of LHb-projecting neurons in the rEPN (p<0.0001, repeated measures two-way ANOVA) (**Fig. 1B, D**). Footshock also increased cFos in LHb-projecting neurons in the LH, which interestingly were located in the ventrolateral portion of the lateral hypothalamus (vlLH), adjacent to the rEPN, but not in the dorsomedial portion (dmLH) (p<0.0001 and p=0.127 for vlLH and dmLH, repeated measures two-way ANOVA) (**Fig. 1C, D**). However, the proportion of CTb-labeled cells expressing cFos in the vlLH was much less than the rEPN (p<0.0001, repeated measures two-way ANOVA). We did not see increased cFos levels in CTB-labeled cells in the other regions examined.

**Fig 1.**
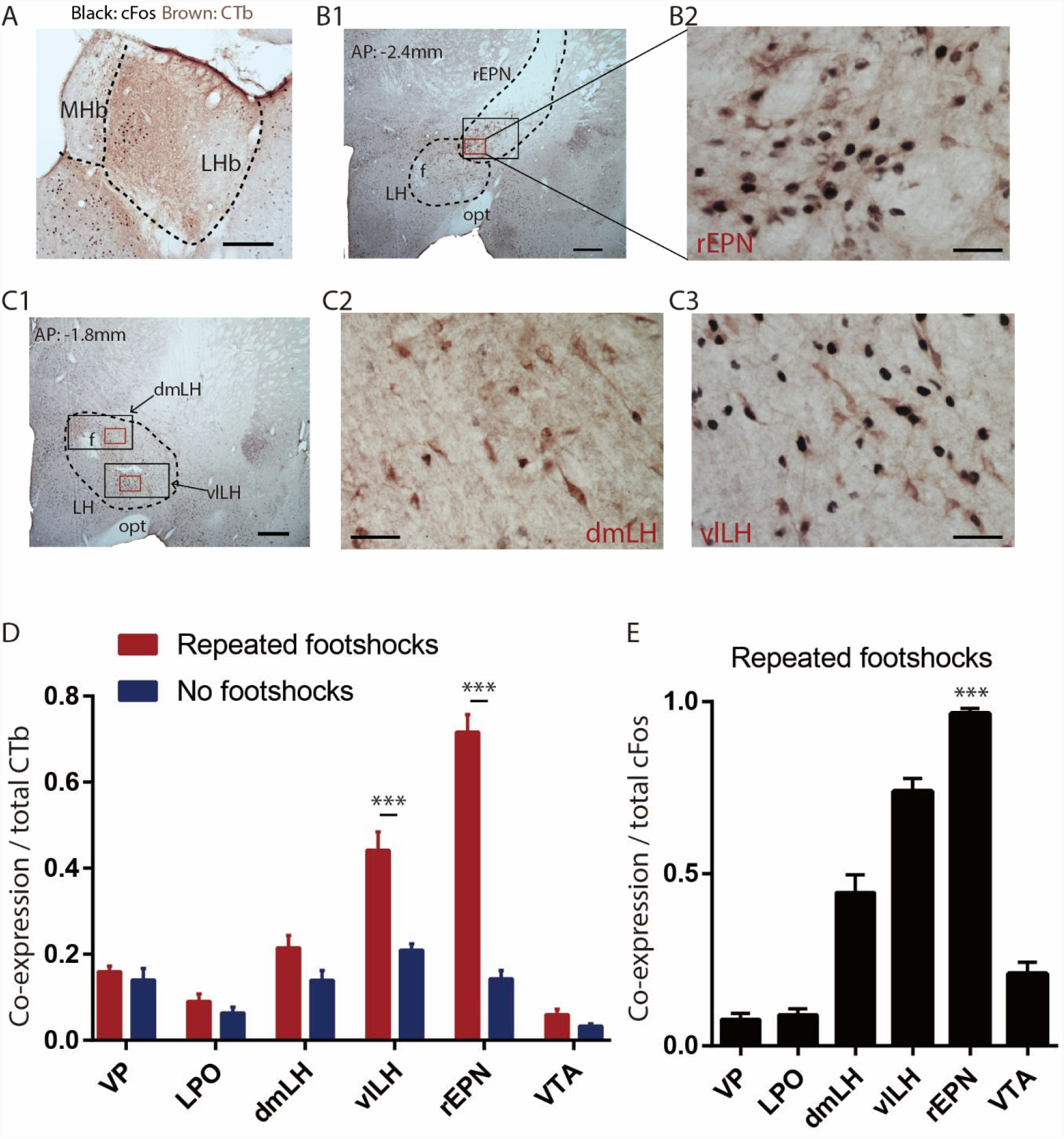
LHb-projecting neurons in the rEPN and the LH are significantly activated by repeated footshocks. Representative photos of immunostaining of CTb (brown) injection site in the LHb. (**B**) Representative photos of immunostaining of CTb (brown) and shock-induced cFos (black) in the rEPN and the LH (**C**). Black squares: areas for cFos counting. Red squares: areas enlarged in representative images. f: fornix; opt: optic tract; MHb: medial habenula. (**D**) Proportion of CTb-labeled neurons expressing cFos in the VTA, rEPN, dmLH, vlLH, LPO, and VP, showing the largest shock-induced increases in the rEPN and LH. (**E**) Proportion of cFos neurons expressing CTb in these same regions. Scalebars 250μm (**A**), 500μm (**B**, **C1**), 100μm (**B2**, **C2** and **C3**).

Furthermore, we observed that not only were most LHb-projecting neurons in the rEPN excited by footshocks, but conversely 97% of aversion-excited neurons in the rEPN were in turn LHb-projecting, again with this percentage higher than in the other regions examined (p=0.01 and p<0.0001 compared to the vlLH and to the remaining regions, respectively, one-way ANOVA) (**Fig. 1E**).

### Reward cue-inhibited rEPN neurons encode valence

Given the large proportion of neurons in the rEPN activated by footshock, we further examined rEPN electrophysiological responses to motivational stimuli in a Pavlovian conditioning paradigm. In this paradigm, three distinct 75dB auditory tones (lasting for 2 seconds) signaled sucrose pellet delivery, brief footshock (0.6 mA, 10-30ms duration), or nothing (**Fig. 2A**). In reward trials, reward delivery was omitted randomly in 20% of trials, in order to examine rEPN responses to negative reward prediction errors. Among 134 recorded neurons, we identified 53 that were from electrodes localized to the rEPN as identified by a lack of parvalbumin immunostaining (Rajakumar et al., 1994), and presence of a high density of retrogradely labeled neurons after injections of the retrograde tracer CTb into the LHb (**Fig. 2B**). Hence, although we did not conclusively identify the projection targets of our neurons, a high proportion are likely to be LHb-projecting. A previous study had further shown that optogenetically identified LHb-projecting EPN neurons in mice exhibit significant inhibition to reward cues between 100-500ms post-stimulus, while EPN neurons projecting to other targets show excitation or no response to reward cues (Stephenson-Jones et al., 2016). Although that study was conducted in mice, previous studies have suggested that anatomic and molecular profiles of rEPN neurons are conserved between rats and mice (Li et al., 2011; Meye et al., 2016; Shabel et al., 2012; Stephenson-Jones et al., 2016), and we further observed that a high proportion of our recorded rEPN neurons also showed inhibition by reward cues 100-500ms post-stimulus (24/53 recorded neurons) (p<0.05, Wilcoxon rank-sum test) (**Fig. 2C**). Hence, we focused our subsequent analyses on these reward cue-inhibited neurons.

**Fig 2.**
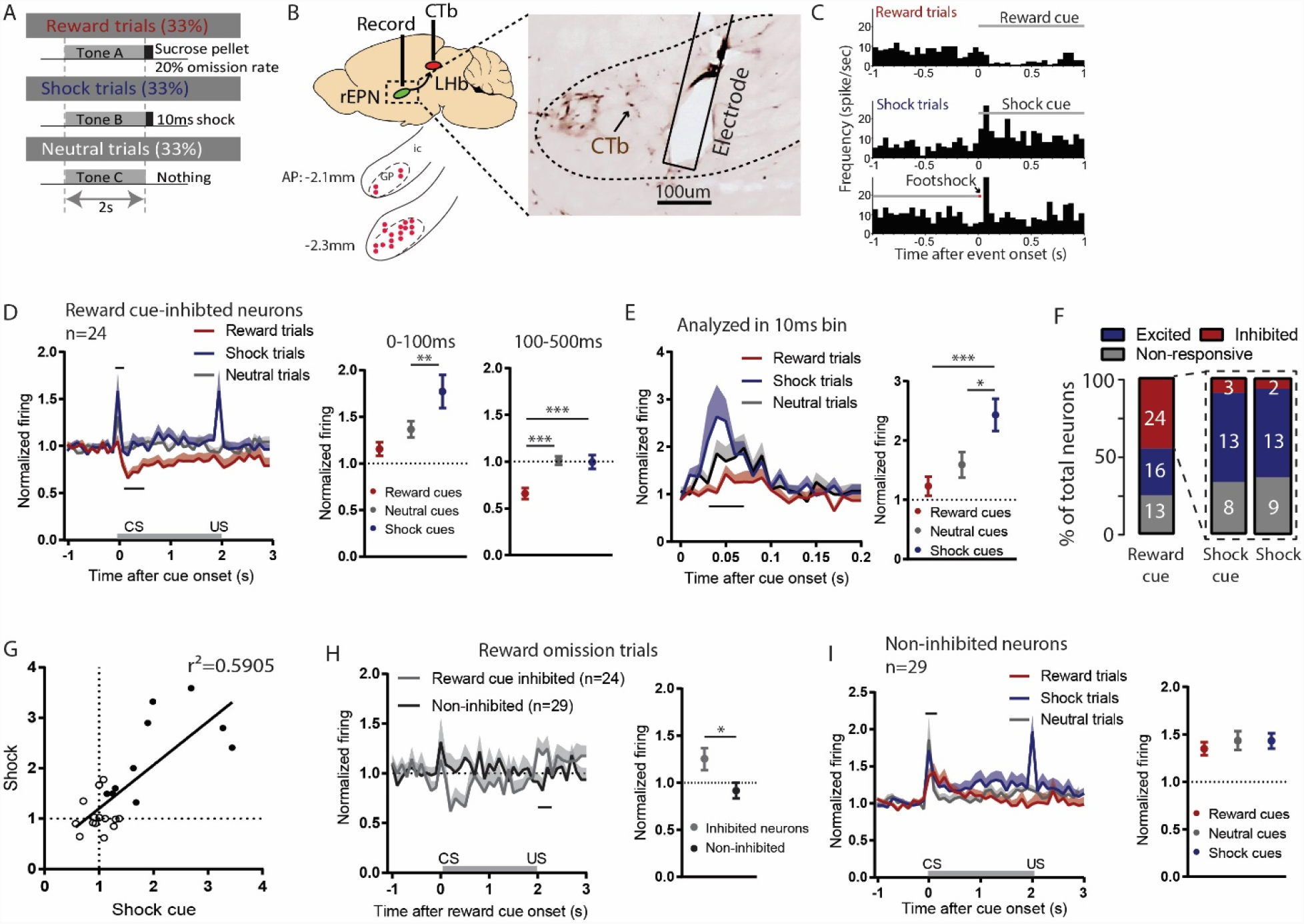
Reward cue-inhibited rEPN neurons encode valence. Schematic of recording paradigm, in which three different auditory tones signaled a sucrose pellet delivery, a mild footshock or nothing. (**B**) We recorded in the rEPN, as delineated by retrograde labeling of CTb from the LHb and absence of parvalbumin immunoreactivity. (**C**) Representative histograms of rEPN firing in response to reward and shock trials, analyzed in 50ms bins. (**D**) Responses of reward cue-inhibited subpopulation to reward, shock, and neutral trials, again in 50ms bins. Excitation to shock cue is most prominent in first two bins (0-100ms, upper solid black line), while inhibition to reward cue is most prominent 100-500ms after stimulus onset (lower solid black line). Average firing rates from these two windows are shown in adjacent graph. (**E**) rEPN response histogram to cues analyzed in 10ms bins, and average firing (30-70ms analysis window). Baseline firings were the same in **D** (not shown). (**F**) rEPN neurons showed heterogeneous responses to reward and shock trials, with predominant inhibition to reward cues and excitation to footshocks and shock cues. (**G**) In reward cue-inhibited rEPN neurons, responses to shock cues were positively correlated with individual’s responses to footshocks. Solid circles indicate neurons significantly excited by both shock cues and footshocks. (**H**) Reward cue-inhibited neurons also showed excitation to reward omission in response histogram and averages (0-200ms window) (**I**) In contrast to reward cue-inhibited neurons, reward cue non-inhibited neurons did not distinguish between cues indicating reward, shock, or neutral outcomes in histogram or averages (0-100ms window).

The population average of reward-cue inhibited neurons showed significant excitations to shock cues and footshocks between 0-100ms post-stimulus when responses were most prominent (z=2.57 and z=2.66, respectively) (**Fig. 2C, D**). These neurons also showed significant excitation to neutral cues (z=1.98), but the magnitude of the excitation was significantly less than to shock cues (p=0.0037, repeated measures one-way ANOVA) (**Fig. 2D**). Since rEPN activations occurred rapidly (within 100ms post-stimulus), we further analyzed data in 10ms bins in order to examine more precisely the temporal characteristics of this excitation. We found that excitations to both shock cues and neutral cues were particularly prominent 30-70ms post-stimulus (z=1.42, z=2.77, and z=5.18 for reward cue, neutral cue, and shock cue, respectively), and that magnitude of shock cue responses during this period were also significantly greater than neutral cues (p=0.0005 and p=0.03 compared with reward cue and neutral cue, one-way ANOVA) (**Fig. 2E**).

When individual neuron responses were analyzed, we found that more than half of reward cue-inhibited neurons showed significant excitation to shock cues (13/24) and footshocks (13/24). A smaller proportion showed either no response to shock cues (8/24) or footshocks (9/24), while the smallest proportion showed significant inhibition (3/24 and 2/24 for shock cues and footshocks) (p<0.05, Wilcoxon rank-sum test) (**Fig. 2F**). Most shock cue-excited neurons (10/13) also exhibited significant excitation to footshock outcomes, and these responses were correlated with each other (r^2^=0.5905, p<0.0001) (**Fig. 2G**). Additionally, we found that reward-cue inhibited neurons were more strongly activated by reward omission compared with non-inhibited neurons (p=0.024, unpaired t-test) (**Fig. 2H**). Together, our results indicate that reward-cue inhibited neurons are most likely to also be activated by shock cues, footshocks, and reward omission, consistent with a valence-encoding pattern.

Neurons that did *not* exhibit inhibition to reward cues (n=29) showed similar magnitude of excitations to reward cues, neutral cues, and shock cues, suggesting a lack of discrimination between motivational significance in these rEPN neurons (F=0.4958, p>0.05, one-way ANOVA) (**Fig. 2I**).

### LHb neurons display two temporal phases in responses to shock cues

Our cFos and electrophysiology results are consistent with a rEPN role in driving LHb responses to motivational stimuli. To more directly test this hypothesis we recorded LHb responses (**Fig. 2A**), in a separate group before and after inactivating the ipsilateral EPN via micro-infusing a cocktail of GABA_A_/GABA_B_ receptor agonists (0.05nmol muscimol and 0.5nmol baclofen) during the latter half of each recording session (**Fig. 3A,B, 4A**). Prior to rEPN inactivation, we found that large proportions of LHb neurons were inhibited by the reward cue (20/32) between 100-500ms post stimuli, and were activated by shock cues (18/32) and footshocks (15/32) between 0-100ms post stimuli (p<0.05, Wilcoxon rank-sum test) (**Fig. 3C**). This is similar to our observed response patterns in reward cue-inhibited rEPN neurons, and also consistent with primate studies showing that LHb neurons encode negative valence (Hong et al., 2011; Jhou et al., 2013). Although responses to shock cues were positively correlated with individual’s responses to footshocks (r^2^=0.2027, p=0.01) (**Fig. 3D**), we unexpectedly found that only 8/32 LHb neurons were significantly excited by both stimuli, while 10/32 and 6/32 LHb neurons showed significant excitations to only shock cues or footshocks (**Fig. 3E**).

**Fig 3.**
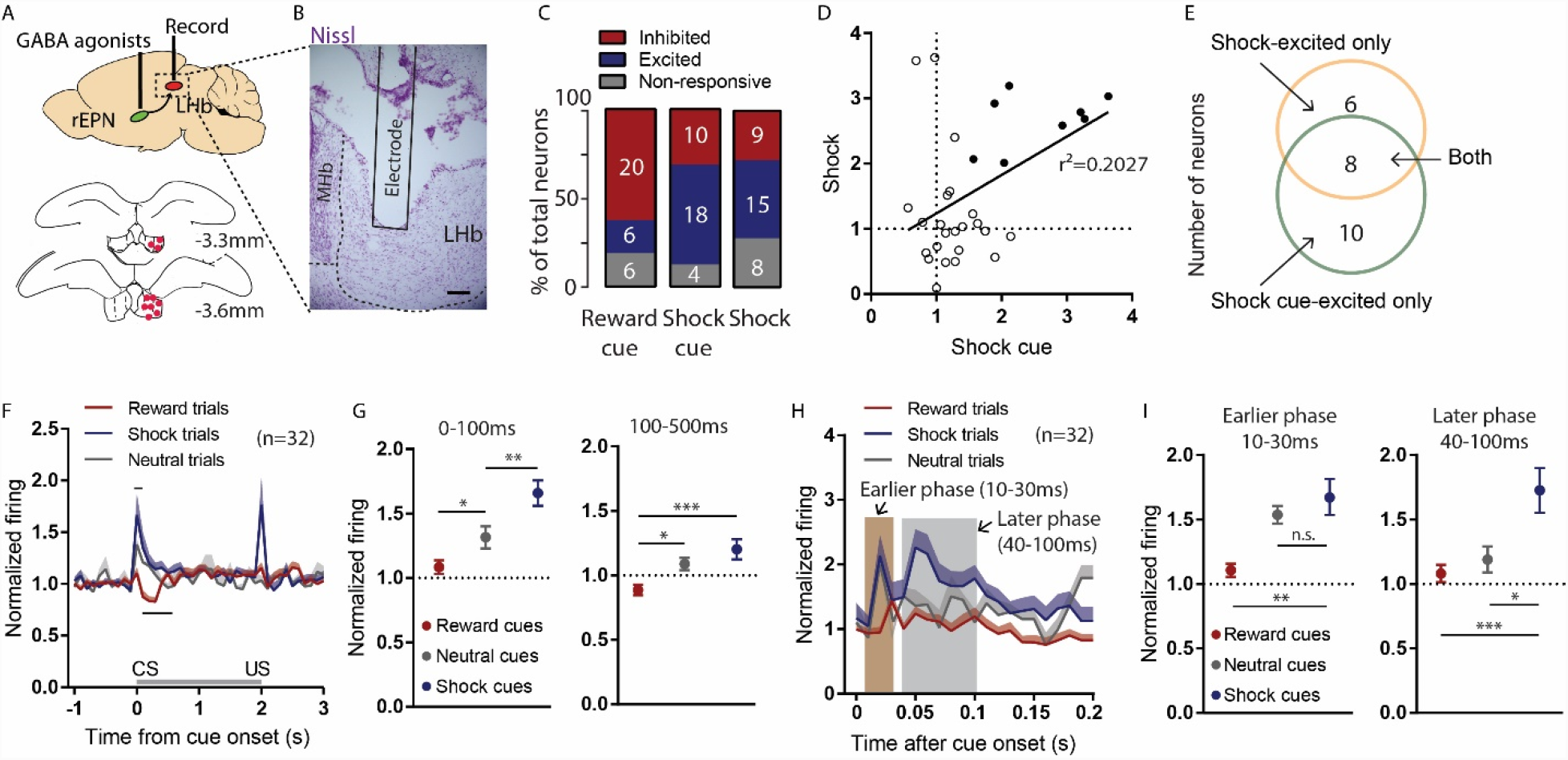
LHb neurons display two temporal phases in responses to shock cues. Schematic of recording from the LHb while inactivating ipsilateral rEPN with GABA agonists. (**B**) Representative photomicrograph of recording electrode track in the LHb. Scalebar: 100um. (**C**) A majority of LHb neurons are inhibited by reward cue and activated by shock and shock cues. (**D, E**) Although responses to shock cues were positively correlated with individual’s responses to footshocks, they were largely encoded in different LHb neurons. Solid circles indicate neurons significantly excited by both shock cues and footshocks. (**F, G**) LHb neurons on average showed slow inhibition to reward cues 100-500ms post-stimulus and rapid excitation to shock cues 0-100ms post-stimulus. Black bars indicate analysis windows for F. (**H, I**) Excitation to shock cues contained two phases: an “earlier” phase from 10-30ms post-stimulus and a “later” phase from 40-100ms post-stimulus. Neutral cues produced a significant response only in the earlier phase, while reward cues did not activate neurons in either phase. Baseline firings were the same in **F** (not shown).

Furthermore, LHb population averages also showed significant inhibition by reward cues during 100-500ms and excitation by both neutral cues and shock cues 0-100ms post-stimulus (z=-28.4, z=2.74, and z=4.813 for reward cues, neutral cues and shock cues, respectively) (**Fig. 3F**). The magnitude of shock cue responses was significantly greater than responses to the neutral cue, similar to responses seen in rEPN neurons (F=13.34, p=0.0085, one-way ANOVA) (**Fig. 3G**). When further analyzing data in 10ms bin, we found that LHb neuron activation to shock cues contained two temporal phases, an “earlier” phase lasting 10-30ms post-stimulus and a “later” phase lasting 40-100ms post-stimulus, while neutral cues only produced an earlier phase activation and reward cues caused no activation (**Fig. 3H**). During the earlier phase, responses to shock cues were not different in magnitude from responses to neutral cues but were significantly greater than responses to reward cues (F=5.58, p=0.275 and p=0.004, respectively, one-way ANOVA). In contrast, during the later response phase, shock cues elicited significantly greater LHb neuron activations than both reward cues and neutral cues (F= 7.829, p=0.0008 and p=0.026, respectively, one-way ANOVA) (**Fig. 3I**). Thus, the two phases of LHb responses potentially encode different information, with the later but not earlier phase responses distinguishing between the aversion-predictive versus neutral cues.

### Temporary inactivation of the rEPN reduces LHb basal activities, eliminates the later phase LHb responses to shock cues, and partially diminishes LHb responses to footshock

After rEPN inactivation, we observed dramatic decreases in basal activities of LHb neurons. Specifically, rEPN inactivation significant reduced LHb basal firing from 3.94 to 2.21 Hz, reduced bursts per minute from 26.26 to 15.8, and reduced percentages of spikes in bursts from 32.5 to 23.97 (t=3.569, p=0.0019; t=4.698, p<0.0001; t=3.621, p=0.0014, respectively, paired t-test) (**Fig. 4B**). rEPN inactivation did not alter mean burst duration of LHb neurons (t=0.5606, p=0.5809, paired t-test) (**Fig. 4C**). These results indicate that LHb basal activities are largely dependent on rEPN inputs.

**Fig 4.**
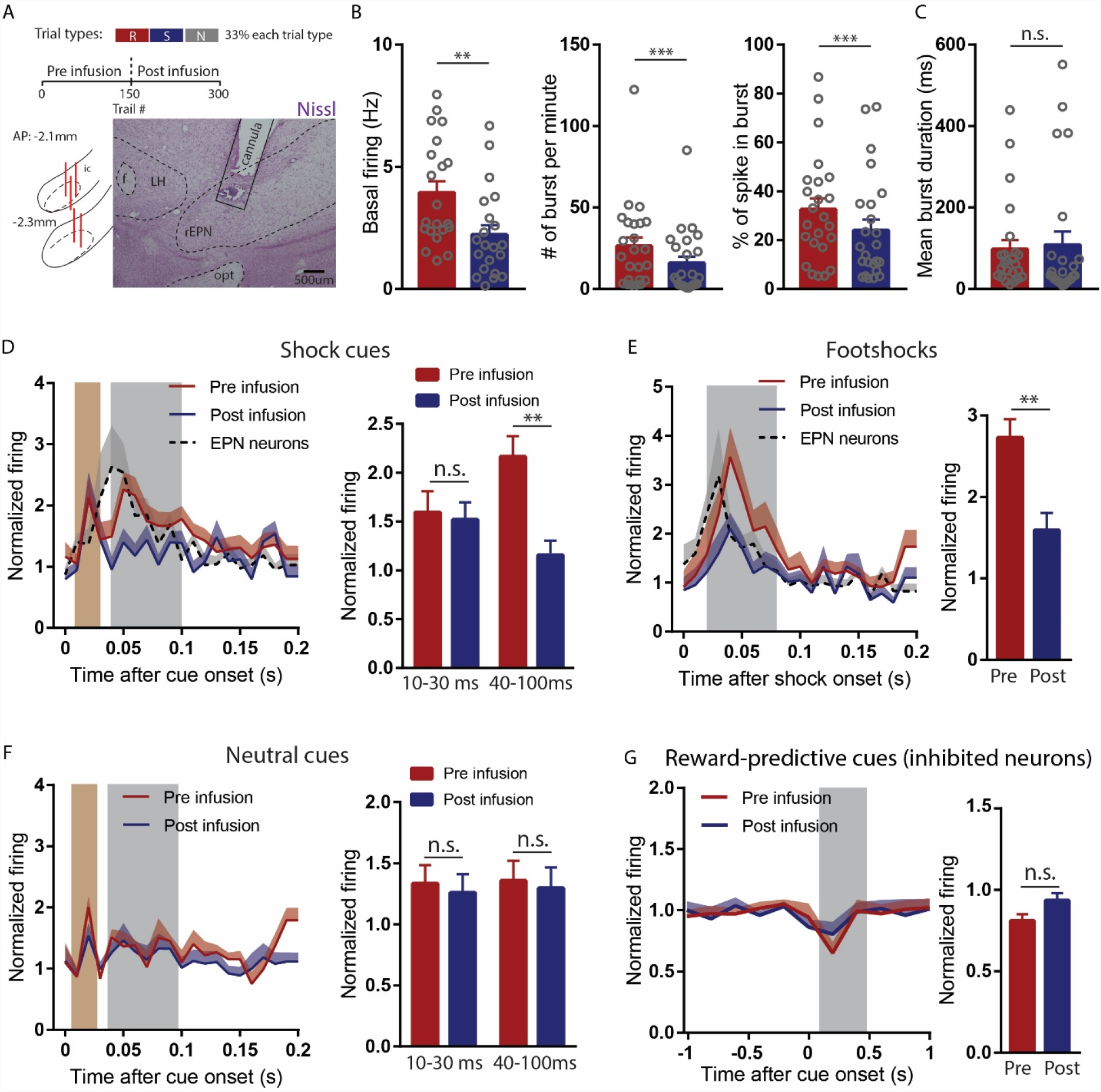
Temporary inactivation of the rEPN reduces LHb basal activities, eliminates the later phase LHb responses to shock cues, and partially diminishes LHb responses to footshock. Schematic of inactivation paradigm and cannula placements in the rEPN. (**B**) rEPN inactivation decreased basal firing, number of burst per minute, and percentage of spikes in bursts in LHb neurons, but not mean burst duration (**C**). (**D**) Ipsilateral rEPN inactivation eliminated later phase LHb responses to shock cues, while having no effects on the earlier phase. (**E**) rEPN inactivation partially reduced LHb reponses to footshocks. (**F, G**) rEPN inactivation had no effects on LHb responses to reward cues nor neutral cues. Baseline firings for **D**, **E**, and **F** were not shown.

In the rEPN recordings above, we had observed activations to shock cues 30-70ms post-stimulus, slightly preceding the later phase LHb response and indicating a potential role of rEPN neurons in driving the later but not earlier phase LHb responses to shock cues. Indeed, after rEPN inactivation, only the later phase LHb responses to shock cues was diminished (F=3.863, p=0.776 and p=0.006 for 10-30ms and 40-100ms, one-way ANOVA), while the earlier phase remained unaffected (**Fig. 4D**). In contrast to the dual response to shock cues, LHb responses to footshocks themselves did not exhibit a clear temporal separation, and most responses occurred 30-70ms post stimuli. Again, rEPN activations to footshocks slightly preceded LHb responses. After rEPN inactivation, LHb responses to footshocks were significantly reduced, but without being entirely eliminated (t=2.953, p=0.007, paired t-test) (**Fig. 4E**). Additionally, LHb responses to either neutral cues or reward cues were unaffected by rEPN inactivation (neutral cue: p>0.05 for both time windows, one-way ANOVA; reward cue: p=0.1, paired t-test) (**Fig. 4F, G**). Thus, our results indicate a selective role for the rEPN in driving LHb responses to negative motivational stimuli, i.e. shocks and shock-predictive cues, but not positive or neutral motivational stimuli.

### Excitotoxic lesions of the rEPN partially reduce footshock-induced cFos in LHb and RMTg

We next tested whether the rEPN was also necessary to drive RMTg activation by footshocks. We unilaterally lesioned the rEPN with an excitotoxin and measured the ratio of ipsilateral over contralateral cFos expression in the LHb and RMTg in response to a series of footshocks (n=4 for lesion group and n=4 for control group). We verified the size and placement of the lesions using NeuN staining (**Fig. 5A, B**), showing greater than 85% loss of rEPN cells in the lesioned side compared with the intact side, with less than 20% cell loss in the adjacent the vlLH region in all cases (t=14.93, p=0.0007, two-tailed paired t-test) (**Fig. 5D**). We found that rEPN lesions significantly diminished footshock-induced cFos on the ipsilateral side of the LHb and the RMTg by 42% and 38% respectively (F=13.68, one-way ANOVA, p=0.0014, p=0.008 for the LHb and the RMTg respectively between lesion and intact). In contrast, sham-lesioned animals showed equal amounts of cFos on both sides (**Fig. 5C, E**). These data suggest that the rEPN plays a partial role in driving LHb and RMTg responses to footshocks.

**Fig 5.**
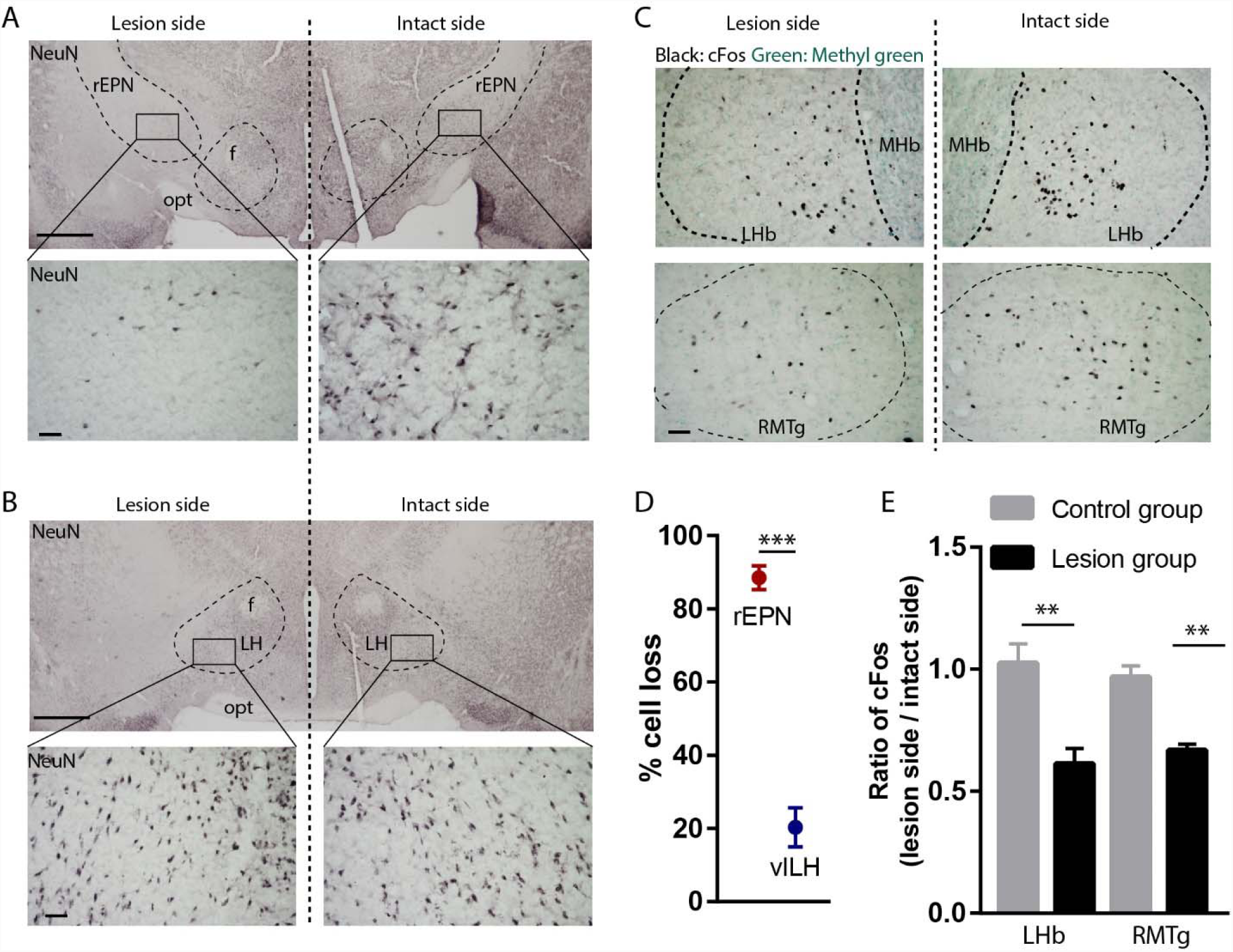
Excitotoxic lesion of the rEPN reduces cFos induced by unsignaled footshocks in LHb and RMTg. (**A, B**) Representative photomicrographs of the ipsilateral (lesioned) and contralateral (intact) rEPN and vlLH immunostained for the neuronal marker NeuN (black label). (**C**) Representative photographs of footshock-induced cFos (black label) in the LHb and RMTg with methyl green counterstain. (**D**) Number of EPN cells dramatically decreased in the lesioned side, while cells in the vlLH were less affected by the lesions. (**E**) rEPN lesion reduced cFos expression induced by unsignaled footshocks in the lesioned side LHb and RMTg, compared with intact side. Scalebars are 1mm in top panels for **A**, **B**, and 100μm in all other panels.

## Discussion

In the present study, we found that the rEPN contributes to specific portions of the LHb response to negative motivational stimuli instead of driving the full profile of LHb responses. In particular, the LHb activation to shock cues show two temporal phases, with the later but not earlier phase being dependent on the rEPN. LHb activations to shocks show only a single temporal phase, which is partially dependent on the rEPN, while LHb responses to non-aversive stimuli (i.e. reward-related or neutral) are not affected at all by rEPN inactivation. Furthermore, we find that rEPN lesions partially reduced footshock-induced cFos in the LHb and RMTg, while rEPN inactivation also reduces basal LHb activities including basal firing and bursting firing. Hence, our findings indicate an important role of the rEPN in driving habenular basal activities and responses to negative stimuli but not neutral or reward-related stimuli.

One striking result from the current study is the finding that the rEPN influence on the LHb is asymmetric, as it contributes to major elements of the LHb responses to shock-predictive cues and footshocks, but not to reward-predictive cues. This may help shed light on a recent study showing that either activating or inhibiting the rEPN-LHb pathway is sufficient to discourage or encourage reinforcement learning in a decision-making task (Stephenson-Jones et al., 2016). In that study, the authors used a task in which animals performing nosepokes at two ports received either a reward or non-reward (omission). That study did not distinguish whether the rEPN manipulations affected the processing of the rewards or of the omissions, but our findings suggest the latter is more likely.

One unexpected finding in the current study was that LHb activation to shock cues was rapid and contained two temporal phases, an earlier phase (10-30ms) and a later phase (40-100ms post-stimulus) that encode different information. The later phase LHb response was present in responses to shock cues, but absent in responses to neutral cues and reward cues, suggesting that the later phase response is related to a cue’s association with negative outcomes. Although several studies had indicated an LHb role in responses to stimuli that predict negative stimuli (Bromberg-Martin and Hikosaka, 2011; Hong et al., 2011; Wang et al., 2017), our findings showed that LHb responses to conditioned stimulus contained distinct temporal phases, of which only the later phase encoded prediction of negative stimuli. Furthermore, we showed that rEPN inactivation selectively eliminated LHb responses during this later phase without affecting the earlier phase, suggesting that these two phases are functionally independent and that the rEPN is critical in driving LHb responses to prediction of negative stimuli. We had recently observed a related type of dual response in RMTg neurons to shock cues, and that the later phase RMTg activation (also occurring around 40-100ms) was completely dependent on the LHb (H. Li and P.J. Vento, unpublished finding). Therefore, our results highly suggest that not only the rEPN drives the later phase LHb responses to shock cues, but also in turn mediates RMTg responses during the same phase, consistent with studies in primates implying that pallidus inputs drive dopamine responses to stimuli signaling non-reward (Bromberg-Martin et al., 2010).

While the later phase LHb response to shock cues is strongly associated with negative valence, the function of the earlier phase response (10-30ms) is less obvious. We noticed that LHb responses during this phase were largely dependent on the type of the current trial. Specifically, this earlier phase response was only present in responses to non-rewarding cues, i.e. a neutral cue or a shock cue, but absent in responses to reward cues. As the neutral cue is not entirely neutral but assumes a modest negative value due to it also indicating reward omission (Cohen et al., 2012), we speculate that the earlier phase LHb response encodes some components of attention to non-rewarding stimuli, analogous to the early excitations of DA neuron to rewarding stimuli (Matsumoto et al., 2016; Schultz, 2016). However, magnitudes of these responses were not different between shock cues and neutral cues, suggesting that the earlier phase LHb response is insensitive to additional aversively-induced negative shifts in motivational value.

In contrast to the relatively unified rEPN contributions to the later phase LHb responses to shock cues, we found that rEPN inactivation only partially reduced LHb responses to footshocks as measured by either firing or cFos, and similarly, rEPN lesions only partly decreased footshock-induced cFos in the RMTg. One possible explanation is that we observed that a subset of LHb-projecting neurons in the vlLH were also activated by footshocks. Since these were not ablated by rEPN lesions, and presumably were not affected by GABA agonist injections, it is possible that vlLH neurons could potentially drive the remaining LHb-RMTg responses to footshocks. Consistent with this possibility, previous studies have indicated that glutamatergic projection from the LH to LHb is aversive and that LH inactivation diminished LHb activation to footshocks (Lecca et al., 2017; Stamatakis et al., 2016). Furthermore, LHb neurons exhibited more heterogeneities in responses to shock and shock cues compared with reward-cue inhibited EPN neurons, further suggesting that other nuclei could contribute to LHb activation (Faget et al., 2018; Tooley et al., 2018). It is also possible that some RMTg neurons receive separate inputs from other nuclei than the LHb, and respond to footshocks regardless whether the LHb or it afferents are inactivated or not (Brown and Shepard, 2013).

In addition to the rEPN contribution to LHb responses to affective stimuli, the present study also indicates that the rEPN contributes substantially to LHb basal firing and bursting patterns. Numerous studies have indicated the close relationship between LHb basal activities and depressive symptoms. Specifically, patients with depression often exhibit LHb hyperactivity and astrocytes-mediated potentiation in bursting firing (Cui et al., 2018; Lawson et al., 2017; Shumake and Gonzalez-Lima, 2003; Yang et al., 2018). Here, we showed that rEPN inactivation dramatically decreased LHb basal firing rates and reduced bursting firing patterns, suggesting that the rEPN also contributes to burst firing patterns in the LHb, and potentially plays a critical role of the rEPN in regulating depressive symptoms. Consistent with this suggestion, previous studies have shown that rEPN neurons co-release glutamate and GABA onto LHb neurons, and the balance between glutamate and GABA transmissions is essential for driving depressive-like behaviors in mice (Baker et al., 2016; Li et al., 2011; Shabel et al., 2014).

In conclusion, we demonstrate the important role of rEPN neurons in mediating LHb responses to negative but not positive or neutral motivational stimuli. Our study potentially provides novel insights into processing of motivational stimuli, which may be disrupted in many psychiatric disorders.

## Methods

### Animals

All procedures were conducted under the National Institutes of Health Guide for the Care and Use of Laboratory Animals, and all protocols were approved by Medical University of South Carolina Institutional Animal Care and Use Committee. Adult male Sprague Dawley rats weighing 250 to 450 g from Charles River Laboratories were paired housed in standard shoebox cages with food and water provided ad libitum until experiments started. Rats were single housed during all experiments. In total, 38 rats were used for these experiments. 16 rats completed recordings. Of these, 10 rats were used for rEPN recoding, and 6 rats for LHb recordings. 14 underwent CTb cFos experiments with 7 rats for footshock group and 7 rats for control group. 8 rats underwent rEPN lesion cFos experiments with 4 rats for lesion group and 4 rats for control group.

### Surgeries

All surgeries were conducted under aseptic conditions with rats that were under isoflurane (1– 2% at 0.5–1.0 liter/min) anesthesia. Analgesic (ketoprofen, 5mg/kg) was administered subcutaneously immediately after surgery. Rats were given at least 5 days to recover from surgery. For rabies tracing experiments, 300nl of retroAAV-cre (Addgene) was injected ipsilaterally with a glass pipette into the VTA (AP: −5.2mm; ML: 2mm; DV: −8.2mm from dura, 10-degree angle) or the RMTg (AP: −7.4mm; ML: 2.1mm; DV: −7.7mm from dura, 10-degree angle). Flexed-TVA-mcherry and Flexed-RG into the LHb (AP: −3.4mm; ML: 1.5mm; DV: −4.5mm from dura, 10-degree angle) at the same time. 21 days later, EnvA-ΔG-rabies-GFP was injected into the LHb. For recording experiments, custom drivable electrode arrays were implanted above the rEPN (AP: −2.3mm; ML: 2.8mm; DV: −7.1mm from dura) or the LHb (AP: −3.4mm; ML: 1.5mm; DV: −4.5mm from dura, 10-degree angle). For CTb injection, 80nl of CTb was injected ipsilaterally into the LHb. For lesion experiments, 50nl of 400mM quinolinic acid per side was injected into the rEPN (AP: −2.5mm; ML: 3.0mm; DV: −7.5mm from dura). Rats were kept anesthetized with pentobarbital intraperitoneally (55mg/kg) for up to 3 hours’ post-surgery to reduce excitotoxic effect.

### Perfusions and tissue sectioning

Rats used for all experiments were sacrificed with an overdose of isoflurane and perfused transcardially with 10% formalin in 0.1M phosophate buffered saline (PBS), pH 7.4. Brains from electrophysiology experiments had passage of 100μA current before perfusion, allowing electrode tips to be visualized. Brains were removed from the skull, and post-fix in 10% formalin for 24 hours before equilibrated in 20% sucrose solution until sunk. Brains were cut into 40μm sections on a freezing microtome. Sections were stored in phosphate buffered saline with 0.05% sodium azide.

### Immunohistochemistry

Free-floating sections were immunostained for CTb, NeuN or cFos by overnight incubation in goat anti-CTb (List Biological Laboratories, 7032A9, 1: 50,000 dilution), mouse anti-NeuN (Millipore, MAB-377, 1: 5,000 dilution), or rabbit anti-cFos (Millipore, ABE457, 1:1,000 dilution) primary in PBS with 0.25% Triton-X and 0.05% sodium azide. Afterwards, tissue was washed three times in PBS and incubated in biotinylated donkey-anti-goat, anti-mouse or anti-rabbit secondary (1:1000 dilution, Jackson Immunoresearch, West Grove, PA) for 30 min, followed by three 30s rinses in PBS, followed by 1 hour in avidin-biotin complex (Vector). For TH-staining, tissue was then rinsed in sodium acetate buffer (0.1M, pH 7.4), followed by incubation for 5 min in 1% diaminobenzidine (DAB). For cFos and NeuN staining, nickel and hydrogen peroxide (Vector) were added to reveal a blue-black reaction product.

### Behavioral training for electrophysiological recordings

Rats were food restricted to 85% of their *ad libitum* body weight and trained to associate distinct auditory cues with either a sucrose pellet or no outcome. Behavior was conducted in standard Med Associates chambers (St. Albans, VT). Reward and neutral cues were a 1 kHz tone (75 dB) and white noise (75dB), respectively. The Reward cue was presented for 2s, and a sucrose pellet (45 mg, BioServ) was delivered immediately after cue offset. The neutral cue was also presented for 2s, but no sucrose pellet was delivered. The two trial types were randomly presented with a 30s interval between successive trials. A “correct” response was scored if the animal either entered the food tray within 2 s after reward cues, or withheld a response for 2 s after neutral cues. Rats were trained with 100 trials per session, one session per day, until they achieved 85% accuracy in any 20-trial block. Once 85% accuracy was established, rats underwent surgeries. After recovery from surgeries, rats were then trained with one extra session in which neutral cue trials were replaced by aversive trials consisting of a 2s 8kHz tone (75dB) followed by a 10ms 0.6mA footshock.

### Electrophysiological recordings

After final training, electrodes consisted of a bundle of sixteen 18 μm Formvar-insulated nichrome wires (A-M system) attached to a custom-machined circuit board. Electrodes were grounded through a 37-gauge wire attached to a gold-plated pin (Newark Electronics), which was implanted into the overlying cortex. Recordings were performed during once-daily sessions, and electrodes were advanced 80-160 μm at the end of each session. The recording apparatus consisted of a unity gain headstage (Neurosys LLC) whose output was fed to preamplifiers with high-pass and low-pass filter cutoffs of 300 Hz and 6 kHz, respectively. Analog signals were converted to 18-bit values at a frequency of 15.625 kHz using a PCI card (National Instruments) controlled by customized acquisition software (Neurosys LLC). Spikes were initially detected via thresholding to remove signals less than twofold above background noise levels, and signals were further processed using principal component analysis performed by NeuroSorter software. Spikes were accepted only if they had a refractory period, determined by <0.2% of spikes occurring within 1ms of a previous spike, as well as by the presence of a large central notch in the auto-correlogram. Neurons that had significant drifts in firing rates and that showed 10% or above similarity in cross-correlogram were excluded. Since the shock duration used in the present study was 10ms, the first 10ms of data after footshock were removed in order to reduce shock artifacts.

For Pavlovian conditioning paradigm, once rats achieved 85% accuracy in reward trials, they were trained to respond to an 8kHz tone (75dB) lasting for 2s followed by a mild footshock (0.6mA). During testing, rats again placed on mild food deprivation, and recordings obtained in sessions consisting of 150 mixture of reward trials, neutral trials, and shock trials randomly selected. Rats were recorded for one or two session per day, and electrodes advanced 80–160 μm at the end of each session. Neurons with significant reductions in baseline firing rates across sessions were excluded from the study, as this is indicative of drifting of recording wires between sessions.

### cFos shock induction

All animals were habituated in the behavioral chamber 20 minutes for 3 days. In the last day, animals received 30 footshocks lasting 5-15 seconds (0.7mA). The induction process lasted 20 minutes. Animals were perfused 1 hour after the end of cFos induction program.

### Statistical analysis of electrophysiological and behavioral data

Burst analysis was performed with NeuroExplorer, with a burst defined by any clusters of spikes beginning with a maximal inter-spike interval of 20 ms and ending with a maximal inter-spike interval of 100 ms. The minimum intra-burst interval was set at 100 ms and the minimum number of spikes in a burst was set at 2. Burst per minute, the percentage of spike firing in bursts, and mean duration of bursts were analyzed.

Neurons with large drifting of the microwire electrodes during recordings were excluded from further analysis. Electrophysiology data were first tested for normality, then transformed to ranked forms if data failed tests of normality (p < 0.05, D’Agostino-Pearson test). Significant responses in neural firing were determined by a threshold of p<0.05 for each neuron’s firing rate versus baseline (Wilcoxon signed-rank). Data were tested for normality (p > 0.05, D’Agostino-Pearson test), and were analyzed using parametric tests. One-way or two-way ANOVA with Holm-Sidak correction for multiple comparisons, and two-tailed t-test were used to compare across experimental conditions, respectively, if not otherwise specified. Calculations were performed using Matlab (Mathworks) and Prism 7 software (Graph Pad).

---

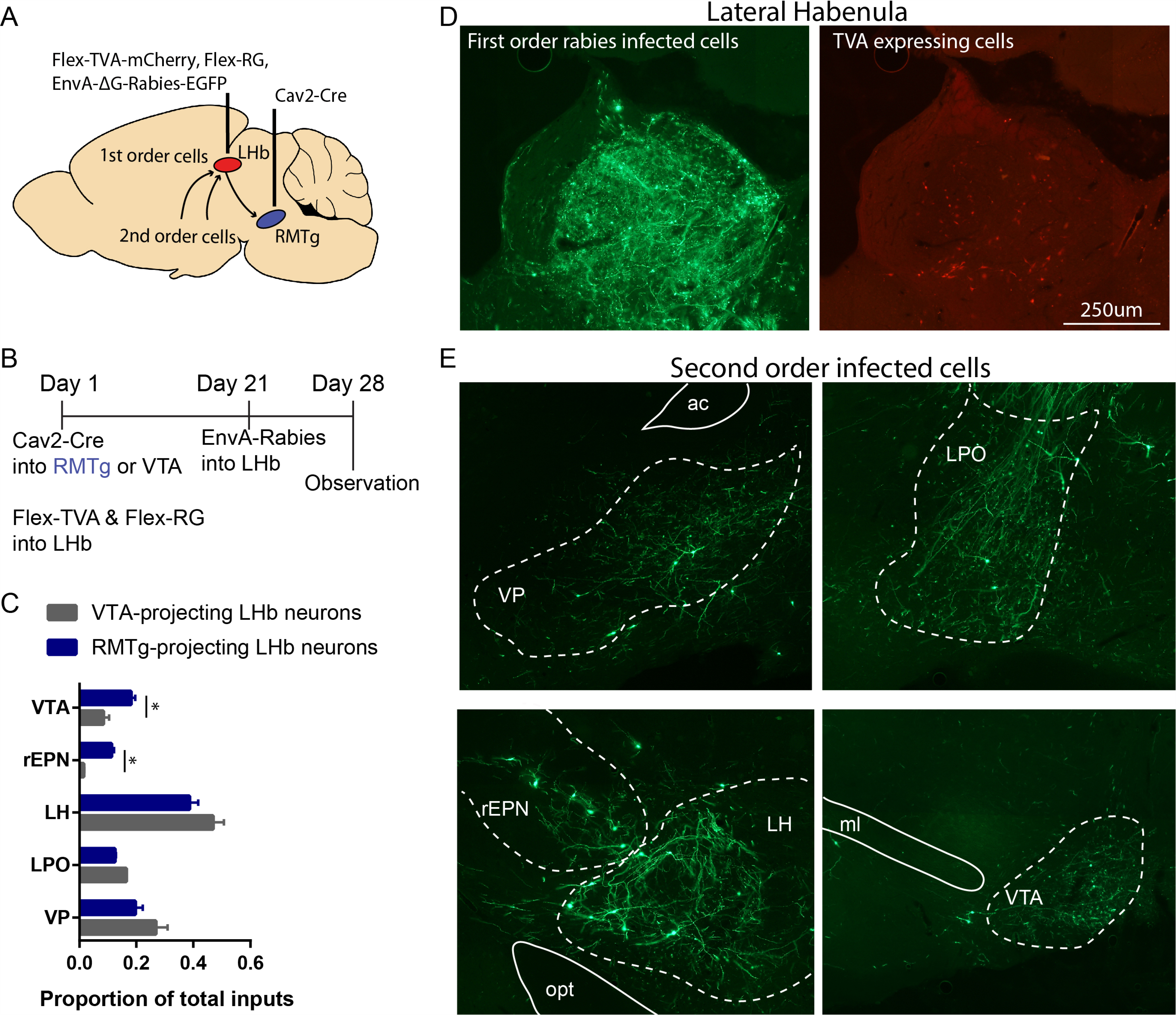

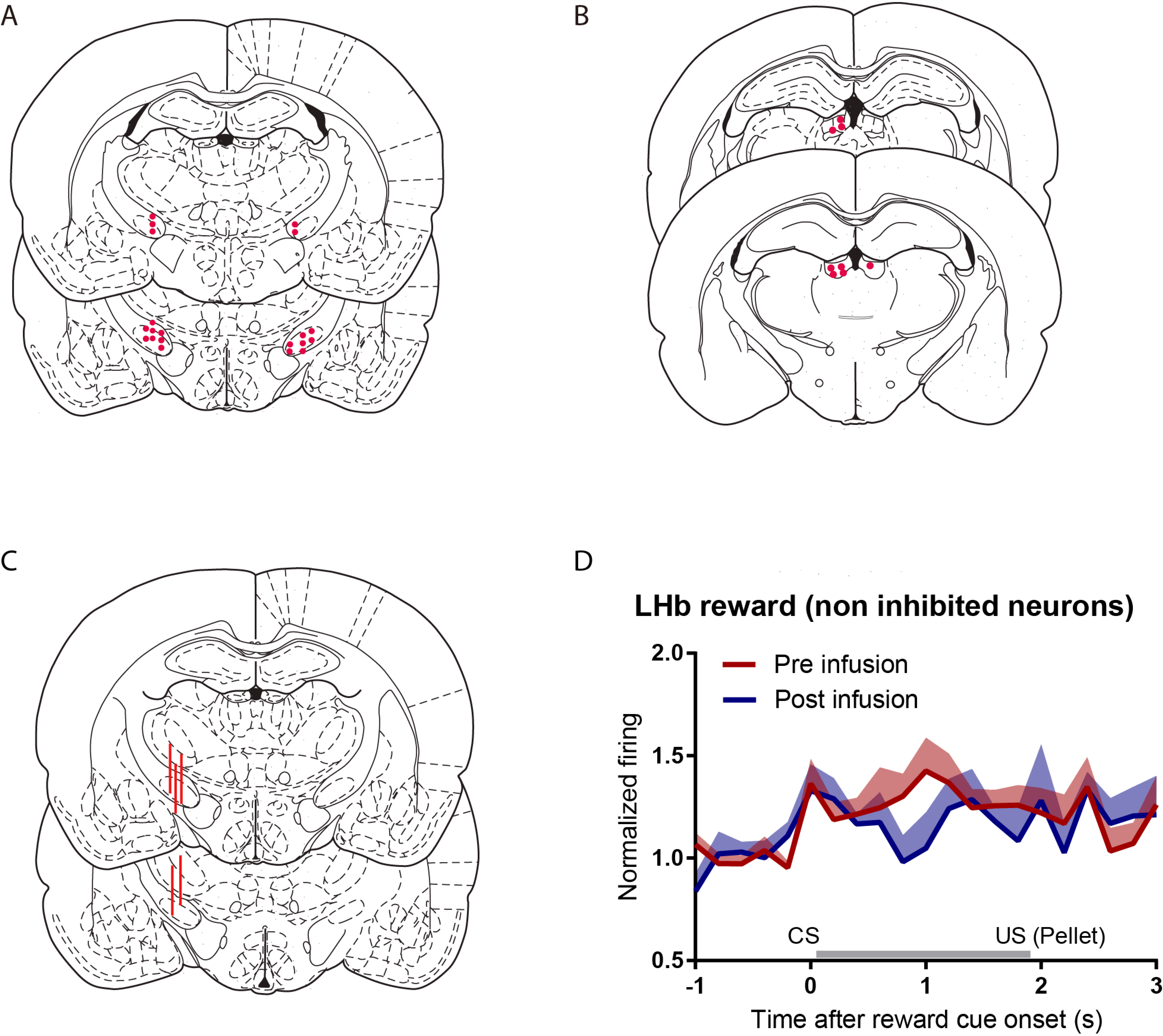

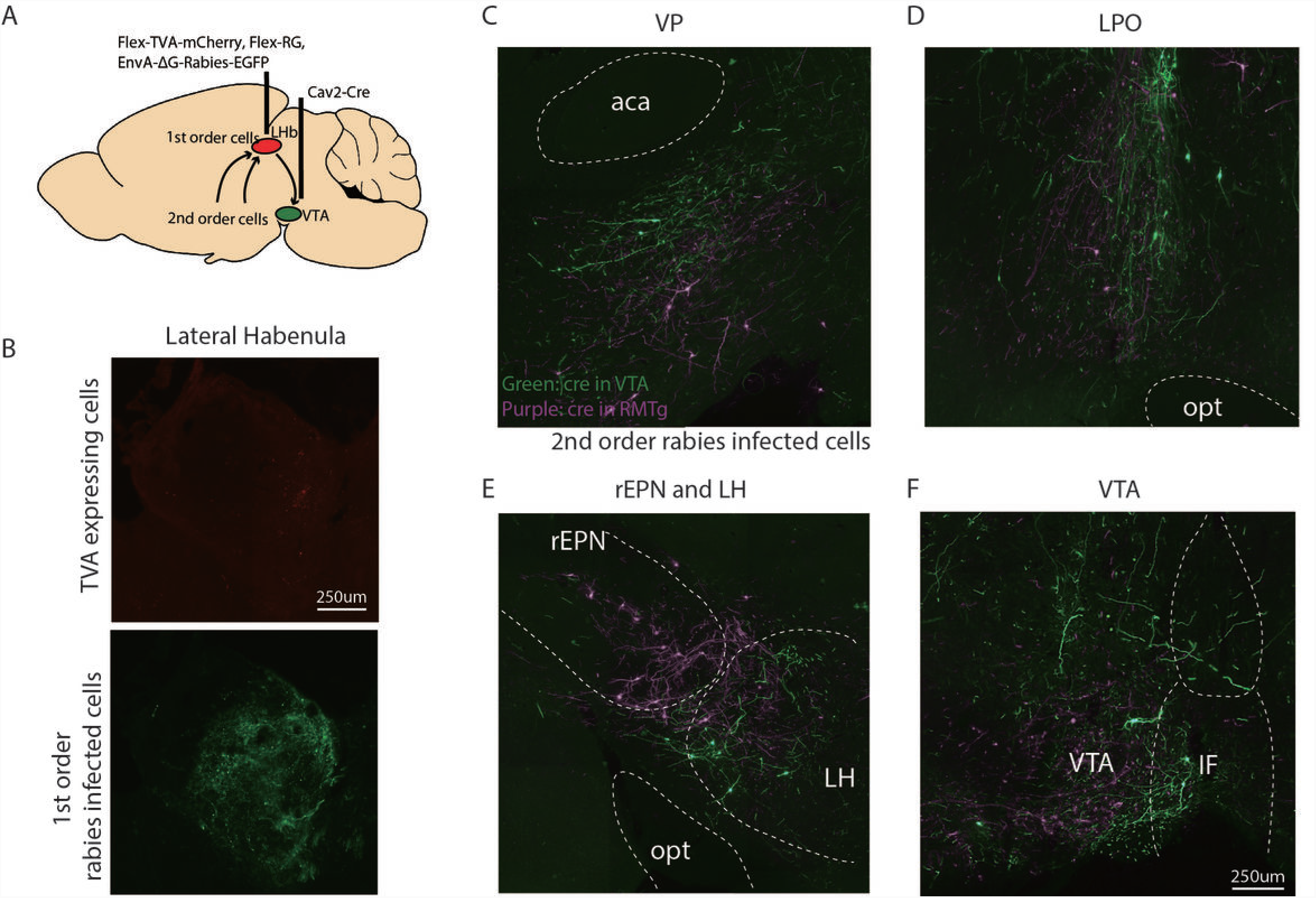

## References

Baker, P.M., Jhou, T., Li, B., Matsumoto, M., Mizumori, S.J., Stephenson-Jones, M., and Vicentic, A. (2016). The Lateral Habenula Circuitry: Reward Processing and Cognitive Control. J Neurosci 36, 11482– 11488.

Bromberg-Martin, E.S., and Hikosaka, O. (2011). Lateral habenula neurons signal errors in the prediction of reward information. Nat Neurosci 14, 1209–1216.

Bromberg-Martin, E.S., Matsumoto, M., Hong, S., and Hikosaka, O. (2010). A pallidus-habenula-dopamine pathway signals inferred stimulus values. J Neurophysiol 104, 1068–1076.

Brown, P.L., Palacorolla, H., Brady, D., Riegger, K., Elmer, G.I., and Shepard, P.D. (2017). Habenula-Induced Inhibition of Midbrain Dopamine Neurons Is Diminished by Lesions of the Rostromedial Tegmental Nucleus. J Neurosci 37, 217–225.

Brown, P.L., and Shepard, P.D. (2013). Lesions of the fasciculus retroflexus alter footshock-induced cFos expression in the mesopontine rostromedial tegmental area of rats. PloS One 8, e60678.

Cohen, J.Y., Haesler, S., Vong, L., Lowell, B.B., and Uchida, N. (2012). Neuron-type-specific signals for reward and punishment in the ventral tegmental area. Nature 482, 85–88.

Cui, Y., Yang, Y., Ni, Z., Dong, Y., Cai, G., Foncelle, A., Ma, S., Sang, K., Tang, S., Li, Y., et al. (2018). Astroglial Kir4.1 in the lateral habenula drives neuronal bursts in depression. Nature 554, 323–327.

Elmer, G.I., Palacorolla, H., Mayo, C.L., Brown, P.L., Jhou, T.C., Brady, D., and Shepard, P.D. (2018). The rostromedial tegmental nucleus modulates the development of stress-induced helpless behavior. Behav Brain Res.

Faget, L., Zell, V., Souter, E., McPherson, A., Ressler, R., Gutierrez-Reed, N., Yoo, J.H., Dulcis, D., and Hnasko, T.S. (2018). Opponent control of behavioral reinforcement by inhibitory and excitatory projections from the ventral pallidum. Nat Commun 9, 849.

Hong, S., Jhou, T.C., Smith, M., Saleem, K.S., and Hikosaka, O. (2011). Negative reward signals from the lateral habenula to dopamine neurons are mediated by rostromedial tegmental nucleus in primates. J Neurosci 31, 11457–11471.

Jhou, T.C., Fields, H.L., Baxter, M.G., Saper, C.B., and Holland, P.C. (2009a). The rostromedial tegmental nucleus (RMTg), a GABAergic afferent to midbrain dopamine neurons, encodes aversive stimuli and inhibits motor responses. Neuron 61, 786–800.

Jhou, T.C., Geisler, S., Marinelli, M., Degarmo, B.A., and Zahm, D.S. (2009b). The mesopontine rostromedial tegmental nucleus: A structure targeted by the lateral habenula that projects to the ventral tegmental area of Tsai and substantia nigra compacta. J Comp Neurol 513, 566–596.

Jhou, T.C., Good, C.H., Rowley, C.S., Xu, S.P., Wang, H., Burnham, N.W., Hoffman, A.F., Lupica, C.R., and Ikemoto, S. (2013). Cocaine drives aversive conditioning via delayed activation of dopamine-responsive habenular and midbrain pathways. J Neurosci 33, 7501–7512.

Ji, H., and Shepard, P.D. (2007). Lateral habenula stimulation inhibits rat midbrain dopamine neurons through a GABA(A) receptor-mediated mechanism. J Neurosci 27, 6923–6930.

Lawson, R.P., Nord, C.L., Seymour, B., Thomas, D.L., Dayan, P., Pilling, S., and Roiser, J.P. (2017). Disrupted habenula function in major depression. Mol Psychiatry 22, 202–208.

Lecca, S., Meye, F.J., Trusel, M., Tchenio, A., Harris, J., Schwarz, M.K., Burdakov, D., Georges, F., and Mameli, M. (2017). Aversive stimuli drive hypothalamus-to-habenula excitation to promote escape behavior. Elife 6.

Li, B., Piriz, J., Mirrione, M., Chung, C., Proulx, C.D., Schulz, D., Henn, F., and Malinow, R. (2011). Synaptic potentiation onto habenula neurons in the learned helplessness model of depression. Nature 470, 535– 539.

Li, K., Zhou, T., Liao, L., Yang, Z., Wong, C., Henn, F., Malinow, R., Yates, J.R., 3rd, and Hu, H. (2013). betaCaMKII in lateral habenula mediates core symptoms of depression. Science 341, 1016–1020.

Matsumoto, H., Tian, J., Uchida, N., and Watabe-Uchida, M. (2016). Midbrain dopamine neurons signal aversion in a reward-context-dependent manner. Elife 5.

Matsumoto, M., and Hikosaka, O. (2008). Negative motivational control of saccadic eye movement by the lateral habenula. Prog Brain Res 171, 399–402.

Meye, F.J., Soiza-Reilly, M., Smit, T., Diana, M.A., Schwarz, M.K., and Mameli, M. (2016). Shifted pallidal co-release of GABA and glutamate in habenula drives cocaine withdrawal and relapse. Nat Neurosci 19, 1019–1024.

Rajakumar, N., Elisevich, K., and Flumerfelt, B.A. (1994). Parvalbumin-containing GABAergic neurons in the basal ganglia output system of the rat. J Comp Neurol 350, 324–336.

Root, D.H., Mejias-Aponte, C.A., Qi, J., and Morales, M. (2014). Role of glutamatergic projections from ventral tegmental area to lateral habenula in aversive conditioning. J Neurosci 34, 13906–13910.

Salas, R., Baldwin, P., de Biasi, M., and Montague, P.R. (2010). BOLD Responses to Negative Reward Prediction Errors in Human Habenula. Front Hum Neurosci 4, 36.

Schultz, W. (2016). Dopamine reward prediction-error signalling: a two-component response. Nat Rev Neurosci 17, 183–195.

Shabel, S.J., Proulx, C.D., Piriz, J., and Malinow, R. (2014). Mood regulation. GABA/glutamate co-release controls habenula output and is modified by antidepressant treatment. Science 345, 1494–1498.

Shabel, S.J., Proulx, C.D., Trias, A., Murphy, R.T., and Malinow, R. (2012). Input to the lateral habenula from the basal ganglia is excitatory, aversive, and suppressed by serotonin. Neuron 74, 475–481.

Shumake, J., and Gonzalez-Lima, F. (2003). Brain systems underlying susceptibility to helplessness and depression. Behav Cogn Neurosci Rev 2, 198–221.

Stamatakis, A.M., Van Swieten, M., Basiri, M.L., Blair, G.A., Kantak, P., and Stuber, G.D. (2016). Lateral Hypothalamic Area Glutamatergic Neurons and Their Projections to the Lateral Habenula Regulate Feeding and Reward. J Neurosci 36, 302–311.

Stephenson-Jones, M., Yu, K., Ahrens, S., Tucciarone, J.M., van Huijstee, A.N., Mejia, L.A., Penzo, M.A., Tai, L.H., Wilbrecht, L., and Li, B. (2016). A basal ganglia circuit for evaluating action outcomes. Nature 539, 289–293.

Stopper, C.M., and Floresco, S.B. (2014). What’s better for me? Fundamental role for lateral habenula in promoting subjective decision biases. Nat Neurosci 17, 33–35.

Tooley, J., Marconi, L., Alipio, J.B., Matikainen-Ankney, B., Georgiou, P., Kravitz, A.V., and Creed, M.C. (2018). Glutamatergic Ventral Pallidal Neurons Modulate Activity of the Habenula-Tegmental Circuitry and Constrain Reward Seeking. Biol Psychiatry.

Wang, D., Li, Y., Feng, Q., Guo, Q., Zhou, J., and Luo, M. (2017). Learning shapes the aversion and reward responses of lateral habenula neurons. Elife 6.

Yang, Y., Cui, Y., Sang, K., Dong, Y., Ni, Z., Ma, S., and Hu, H. (2018). Ketamine blocks bursting in the lateral habenula to rapidly relieve depression. Nature 554, 317–322.

Yetnikoff, L., Cheng, A.Y., Lavezzi, H.N., Parsley, K.P., and Zahm, D.S. (2015). Sources of input to the rostromedial tegmental nucleus, ventral tegmental area, and lateral habenula compared: A study in rat. J Comp Neurol 523, 2426–2456.

